# Resilience and evolvability of protein-protein interaction networks

**DOI:** 10.1101/2020.07.02.184325

**Authors:** Brennan Klein, Ludvig Holmér, Keith M. Smith, Mackenzie M. Johnson, Anshuman Swain, Laura Stolp, Ashley I. Teufel, April S. Kleppe

## Abstract

Protein-protein interaction (PPI) networks represent complex intra-cellular protein interactions, and the presence or absence of such interactions can lead to biological changes in an organism. Recent network-based approaches have shown that a phenotype’s PPI network’s *resilience* to environmental perturbations is related to its placement in the tree of life; though we still do not know how or why certain intra-cellular factors can bring about this resilience. One such factor is gene expression, which controls the simultaneous presence of proteins for allowed extant interactions and the possibility of novel associations. Here, we explore the influence of gene expression and network properties on a PPI network’s resilience, focusing especially on ribosomal proteins—vital molecular-complexes involved in protein synthesis, which have been extensively and reliably mapped in many species. Using publicly-available data of ribosomal PPIs for *E. coli, S.cerevisae*, and *H. sapiens*, we compute changes in network resilience as new nodes (proteins) are added to the networks under three node addition mechanisms—random, degree-based, and gene-expression-based attachments. By calculating the resilience of the resulting networks, we estimate the effectiveness of these node addition mechanisms. We demonstrate that adding nodes with gene-expression-based preferential attachment (as opposed to random or degree-based) preserves and can increase the original resilience of PPI network. This holds in all three species regardless of their distributions of gene expressions or their network community structure. These findings introduce a general notion of *prospective resilience*, which highlights the key role of network structures in understanding the evolvability of phenotypic traits.

**Author Summary:** Proteins in organismal cells are present at different levels of concentration and interact with other proteins to provide specific functional roles. Accumulating lists of all of these interactions, complex networks of protein interactions become apparent. This allows us to begin asking whether there are network-level mechanisms at play guiding the evolution of biological systems. Here, using this network perspective, we address two important themes in evolutionary biology (i) How are biological systems able to successfully incorporate novelty? (ii) What is the evolutionary role of biological noise in evolutionary novelty? We consider novelty to be the introduction of a new protein, represented as a new “node”, into a network. We simulate incorporation of novel proteins into Protein-Protein Interaction (PPI) networks in different ways and analyse how the resilience of the PPI network alters. We find that novel interactions guided by gene expression (indicative of concentration levels of proteins) creates a more resilient network than either uniformly random interactions or interactions guided solely by the network structure (preferential attachment). Moreover, simulated biological noise in the gene expression increases network resilience. We suggest that biological noise induces novel structure in the PPI network which has the effect of making it more resilient.

## 2 Introduction

The evolutionary mechanisms by which novel information is incorporated in biological systems are not entirely understood. Recently, with the increased availability of genomic data, more attention has been devoted to gaining insight into this phenomenon of evolutionary emergence. Part of the challenge to understand evolutionary emergence is that any introduced feature poses a risk of introducing deleterious effects to a biological system. An extensive amount of research has been dedicated to understand the evolutionary trajectory for protein sequence evolution, and what may enable adaptation without disrupting already present biological functions. Evolutionary emergence has been explored by studying gene duplication [1–3], *de novo* gene emergence [4–7, 40, 53], open reading frame extension [8, 9, 11], and sequence properties [10, 54], i.e. GC-content [12] and codon usage [13, 14]. However, whether a novel protein is deleterious or beneficial depends not only on its own sequence features but also the environmental context of interaction partners [15, 16]. While there has been much focus on addressing wherefrom and how novel sequence features *emerge*, limited attention has been given to how novelty may become *integrated* into the cellular apparatus, from a system-level perspective. One study suggests that the robustness of the composition of protein-protein interaction (PPI) networks may play a role in the successful incorporation of a novel protein into the network [17]. Building on a similar systems-approach in this work, we seek to address which features enable robust integration of novelty on a system-level, by analysing redundancy and perturbations to protein-protein interaction (PPI) networks.

Biological redundancy refers to two or more components performing equivalent functions in a given biological system and that deactivation of one of them has negligible consequences on the performance of the biological phenotype. Previous research has shown that there is a positive association between biological redundancy and network connectivity [18, 19]. In the context of protein networks, biological redundancy has been found to enable robustness and increased tolerance for perturbations [20]. Here, a perturbation is defined as an alteration; either adding or removing a protein of the given network. Maintaining functionality in the face of perturbations is typically referred to as phenotypic robustness in evolutionary biology [21]. Here, we will primarily use the term “network resilience”, following Zitnik et al. (2019), which describes the extent to which random node removal deteriorates network structure. Assuming that tolerance for novelty is linked to network resilience, we aim to analyse which features affect resilience and enable successful integration of novel proteins into protein-protein interaction networks.

Here, we use network science to computationally explore how novel proteins may become integrated in PPIs. Specifically, we introduce and apply a novel network measure referred to as the *prospective resilience*. This involves introducing new proteins to a network based on different attachment rules and measuring the resulting network’s resilience compared to baseline. By measuring the change in network resilience following the addition of new nodes to the network, we are able to infer how robust a given network structure is to incorporating novel proteins. More information about the resilience measure as well as its behaviour in different types of random networks, such as networks generated by preferential attachment, is found in Section 6.1.

Gene expression is observed to be the strongest predictor for evolutionary rate of proteins, and while the underlying causes are being debated [22–25], it has been suggested that network topology and protein abundance (gene expression) are interlinked [26]. For these reasons, we aim to infer whether gene expression influences network topology and resilience. To do so, we compare the prospective resilience of ribosomal PPI networks under three different mechanisms for attaching novel proteins to the network: a gene expression-based attachment mechanism, a random-attachment mechanism, and a degree-based attachment mechanism. Here, we specifically focus on ribosomes as they translate genetic information from mRNA into proteins and are an essential cellular complex present across all domains of life. We make use of publicly available data (STRING [27] and SNAP [28] databases), which are thoroughly annotated and experimentally verified, for three organisms: *Escherichia coli* (prokaryote), *Saccharomyces cerevisiae* (unicellular eukaryote) and *Homo sapiens* (multicellular eukaryote). In the following sections, we define and describe the behavior of this prospective resilience measure, and we outline several new implications of approaching evolvability and resilience from a network perspective.

## 3 Results

### 3.1 Network resilience and prospective resilience

In this section we will introduce *network resilience* and *prospective resilience*, which will motivate the presentation of our results in Section 3.2.1. For full technical details, see Section 5.2. In biological terms, individual nodes represent individual proteins of the PPI network, and we infer the phenotypic robustness by inferring network resilience. The network resilience, *R*, is an information theoretic measure that describes the extent to which random node removal deteriorates network structure [20]. It is computed iteratively, involving the incremental isolation of (i.e. removal of all links to) more and more nodes in the network. In biological terms, links represent protein interactions, and the removal of links represents the removal of an interaction between two proteins, yielding isolated and non-interacting proteins. The number of nodes isolated is the fraction 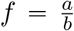 of all nodes in the network (rounded to the nearest number of nodes), where *b* is the total number of iterations and *a* increases from 0 to *b* in steps of 1, i.e, if *b* = 100, we remove 0%, 1%, 2%, … 100% of the nodes. At each iteration, a modified Shannon diversity measure,

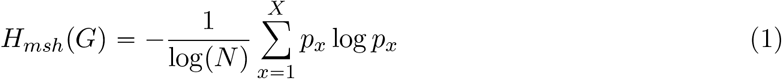

is computed for the resulting network, where 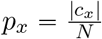 and *c_x_* is a connected component of the network; *p_x_*, therefore, is the probability that a randomly selected node is in the connected component *c_x_*. As *f* increases from 0 to 1, the network becomes more and more disconnected until *f* = 1, at which point the resulting network, *G*_*f*=1_, is a collection of *N* isolated nodes (Figure 1A & 1B). Consequently, the Shannon diversity of these component size distributions increases in *f* (Figure 1C). The final value for resilience is then calculated as a discrete approximation of the area under this curve:

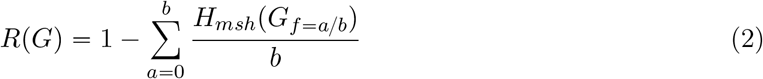

where *H_msh_*(*G_f_*) is the modified Shannon diversity of the network after *f* fraction of nodes have been disconnected. In Supplemental Information (SI) 6.1, we break down the typical behavior of this resilience measure. Particularly, we show that dense Erdős-Rényi networks are more resilient than sparse ones (SI Figure 7), which conforms to the intuition that a complete network is the most resilient network, with a value *R*(*G*) = 0.5. Note that this measure was previously defined as ranging from 0.0 to 1.0 [20], but we show that the theoretical maximum is in fact 0.5 (SI 6.1).

**Figure 1:**
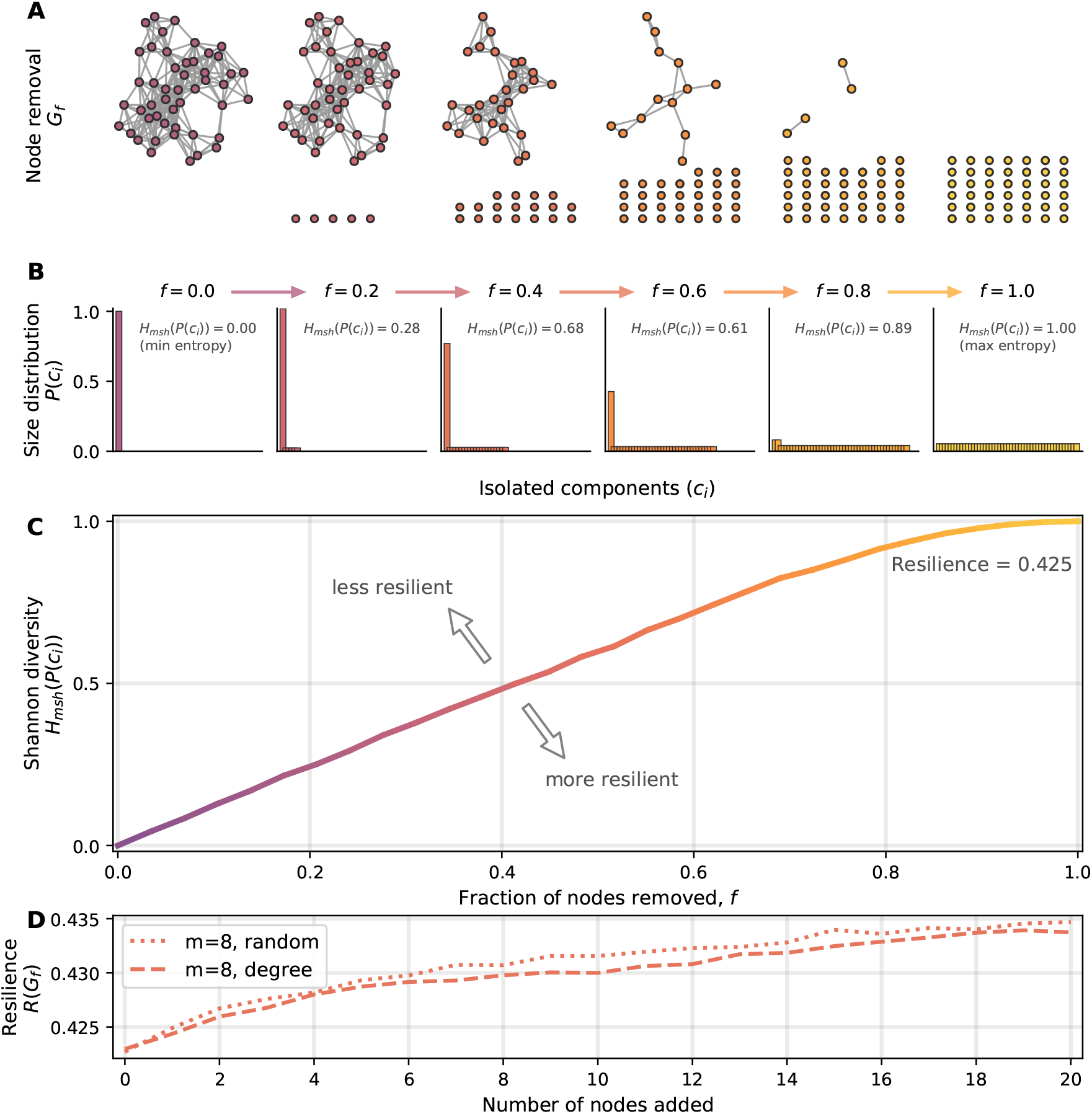
Change in the Shannon diversity indicates network resilience. Here we provide a visual intuition about how network structure is associated with a particular resilience value. **(A)** Network resilience is calculated by iteratively isolating fractions of nodes in the network, *f*, eventually leaving *N* isolated nodes. **(B)** Following every iteration, the Shannon diversity of the component size distribution is calculated, in this case starting at *f* = 0 (one connected component), and increasing until every node is disconnected, *f* = 1. **(C)** Increasing the fraction of nodes that have been removed creates a curve of increasing entropy values, which is used to compute the network resilience, as in Equation 2. **(D)** An example of the *prospective resilience* of the network shown in **(A)**. New nodes are iteratively added to the original network, with links attached randomly or preferentially based on the degree of nodes in the network.

Here, we introduce a novel adaptation of this resilience measure, which we refer to as the prospective resilience (*PR*). The intuition behind this measure is to ask to what extent the resilience of a given network changes following the addition of new nodes into the network structure. In a biological context, this models how a network responds to the introduction of new proteins. Building on common modeling techniques for studying network growth processes, the prospective resilience is obtained by repeatedly adding new nodes to the network and calculating the updated resilience of the resulting network. This yields a vector of resilience values, {*R*_*t*+1_(*G*), *R*_*t*+2_(*G*), …, *R_τ_*(*G*)}, corresponding to the resilience of the network after the addition of each of the *τ* new nodes to the network:

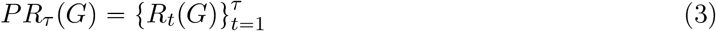

Given that the prospective resilience is computed by adding nodes to a network, the mechanism by which nodes are added becomes an important consideration. In general, node attachment mechanisms assign a probability that each incoming node, *υ*_*t*+1_, attaches its *m* disconnected links (often referred to as “dangling” links) to nodes already in the network, *υ_i_* ∈ *V*. This could be based on *random* attachment, where each node, *υ_i_*, has a probability 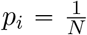 of becoming connected to the incoming node, *υ*_*t*+1_. Similarly, a new node can add its *m* links preferentially based on the *degree* (number of neighbors) of the nodes in the network, 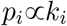, where *k_i_* is the degree of node *υ_i_*. This means that the probability that *υ_i_* will receive an incoming link is 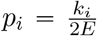, where *E* is the total number of links in the network.

From the biological perspective, we posited that a novel protein entering a system is inevitably more likely to interact with proteins that are more abundant in that system. This abundance can be determined by the protein’s gene expression [29, 30]. To this end, we compare the random and degree-based attachment mechanisms with attachment based on gene expression. This is implemented exactly as for degree-based attachment; the probability that node *υ_i_* receives an incoming link is proportional to *υ_i_*’s gene expression (i.e., the gene expression of node *υ_i_* divided by the sum of the gene expressions of all nodes). New nodes (novel proteins) will not have a known gene expression, and as such, we assign them the average gene expression of the network. Through this attachment rule, we explicitly couple insights from network science to the biological properties of protein networks.

### 3.2 Ribosome protein networks

In this work, we explore the notion of prospective resilience in biological systems. To do so, we focus on ribosomal protein networks: *S. cerevisiae, E. coli*, and *H. sapiens*. In this section, we introduce the procedure for generating these ribosomal protein networks.

Each ribosomal protein of a given species is represented by a node in the network. The links between nodes were then established wherever there was evidence of protein-protein interactions in that species, based on data from the SNAP database [20, 28]. We identified ribosomal proteins from data in [31] and constructed the ribosomal protein networks based on data from the STRING database [27].

Expression for *S. cerevisiae* came from NCBI GEO [32, 33], *H. sapiens* from EMBL-EBI Expression Atlas [34, 35], and *E. coli* K12 from NCBI GEO [32, 36]. See SI 5.1 for a detailed description on how the networks were constructed and how their associated gene expression data was collected. Visualisations of these networks are shown in Figures 2A, 2B, and 2C, and several network properties reported in Table 1. In Figures 2D, 2E, and 2F, distributions of gene expression for each network are plotted as histograms and against node degree. The distributions for all three species had heavy tails, with small numbers of highly expressed proteins and a bulk of proteins with relatively low expression. Across the three networks included here, we see that nodes with similar gene expression and degree tend to cluster together, however the correlation between degree and gene expression itself varies between species (Figures 2G, 2H, and 2I, with Spearman rank correlation coefficients included).

**Figure 2:**
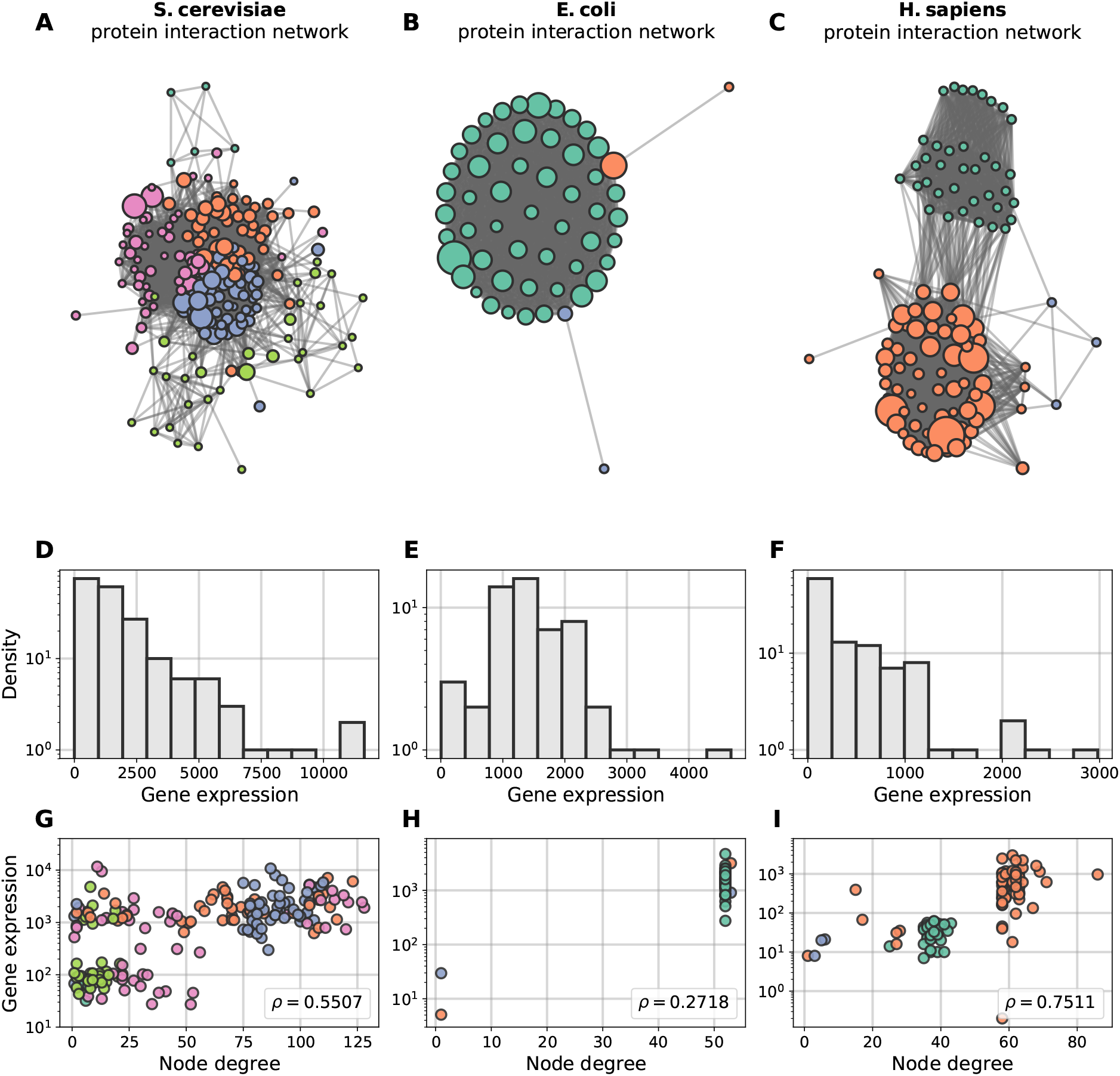
Ribosomal networks of three model species. These species have ribosomal interaction networks that span a range of different network structures. Colours in this plot depict detected communities in the networks—nodes of a given colour are more likely to connect to other nodes of that colour. Node size is proportional to gene expression. **(A)** *S. cerevisiae* ribosomal network. **(B)** *E. coli* ribsomoal network. **(C)** *H. sapiens* ribosomal network. Panels **(D)**, **(E)**, and **(F)** show the gene expression distribution for the three model ribosomal networks discussed in the paper. Panels **(G)**, **(H)**, and **(I)** shows the gene expression against node degree on a scatter plot for the three networks respectively. To accentuate clusters of nodes that share degree and gene expression attributes, the points in these plots share the same color as their corresponding nodes in Figure 2. Of particular note: the gene expression distribution of these three networks are skewed and non-uniform, often referred to as heavy-tailed.

**Figure 3:**
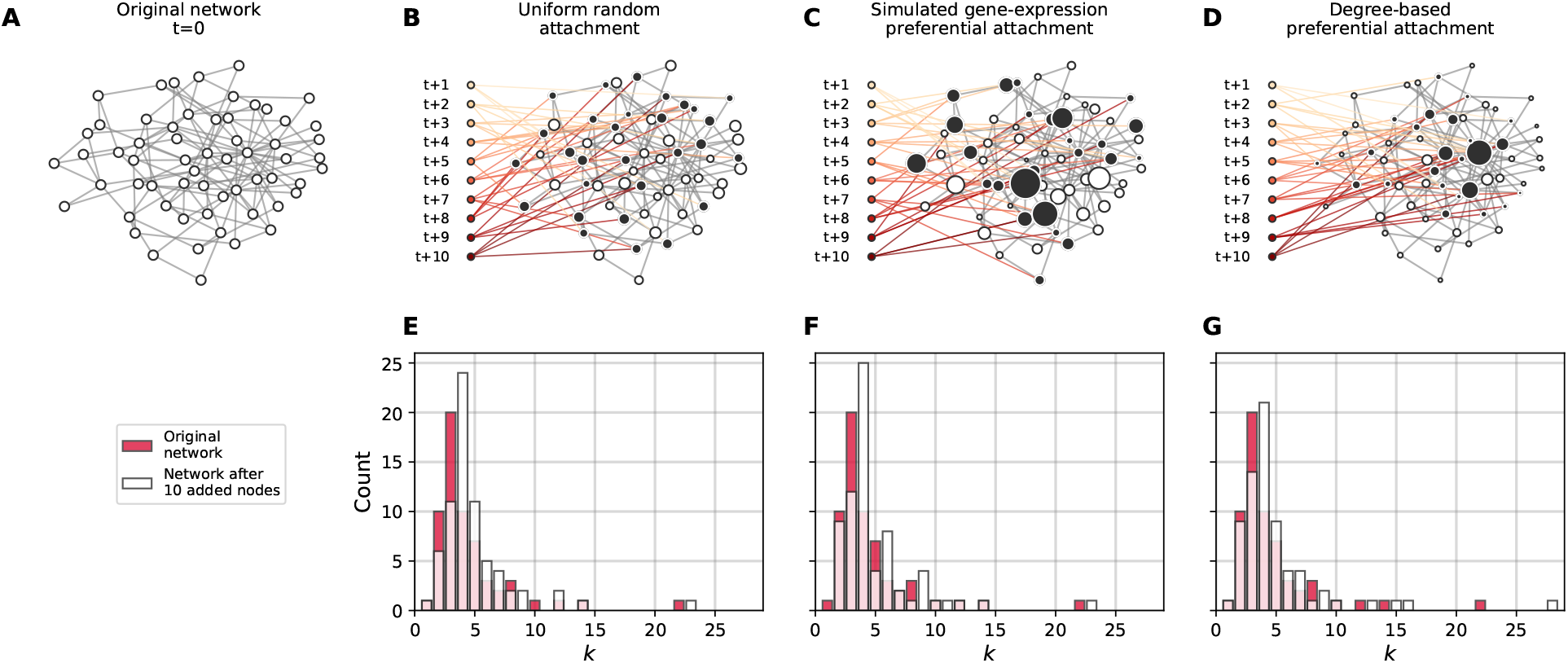
The effect of attachment mechanism on network structure. Here, we offer further intuition about the effect of adding nodes under different attachment mechanisms. In each example, 10 nodes are added, connecting their *m* = 4 links to nodes in the original network (indicated by the black nodes). Node size corresponds to its likelihood of gaining new links. **(A)** Example network, before any new nodes have been added to it. **(B)** Example of uniform attachment. **(C)** Example of (simulated) gene expression preferential attachment. **(D)** Example of degree-based preferential attachment. **(E)—(G)** Histograms showing the change in the original network’s degree distribution after the addition of 10 nodes, under each attachment mechanism. While these histograms highlight the change in a single network property (degree, *k*), one can imagine a number of structural changes occurring following the addition of new nodes, depending on the attachment mechanism.

**Table 1:** Basic network measures of the three networks studied.

| Network property | *S. cerevisiae* | *E. coli* | *H. sapiens* |
| --- | --- | --- | --- |
| Network size | 145 | 55 | 105 |
| Density | 0.284 | 0.929 | 0.471 |
| Average degree | 40.82 | 50.18 | 48.93 |
| Resilience | 0.438 | 0.435 | 0.444 |
| Average clustering | 0.685 | 0.962 | 0.898 |
| Modularity | 0.182 | 0.0013 | 0.363 |

**Table 2:** Spearman rank correlation between the degree and gene expression of a network at different levels of noise. The table displays the correlation after *Noise %* has been introduced to the network. The Spearman correlation was run over the mean from 1000 iterations.

| Species | Noise | $\rho$ | p-value |
| --- | --- | --- | --- |
| *S. cerevisia* | 0.0% | 0.55 | $1.06e^{-16}$ |
| | 20.0% | 0.44 | $1.03e^{-10}$ |
| | 40.0% | 0.33 | $2.18e^{-06}$ |
| | 60.0% | 0.22 | $1.84e^{-03}$ |
| | 80.0% | 0.11 | $1.22e^{-01}$ |
| | 100.0% | -0.0 | $5.17e^{-01}$ |
| *E. coli* | 0.0% | 0.27 | $4.47e^{-02}$ |
| | 20.0% | 0.23 | $4.47e^{-02}$ |
| | 40.0% | 0.16 | $1.72e^{-01}$ |
| | 60.0% | 0.11 | $3.07e^{-01}$ |
| | 80.0% | 0.06 | $4.04e^{-01}$ |
| | 100.0% | -0.0 | $4.78e^{-01}$ |
| *H. sapiens* | 0.0% | 0.75 | $2.78e^{-20}$ |
| | 20.0% | 0.61 | $4.04e^{-12}$ |
| | 40.0% | 0.45 | $1.23e^{-06}$ |
| | 60.0% | 0.31 | $1.47e^{-03}$ |
| | 80.0% | 0.15 | $1.21e^{-01}$ |
| | 100.0% | -0.0 | $4.95e^{-01}$ |

#### 3.2.1 Prospective resilience in ribosomal networks

We here report the prospective resilience of the three ribosomal networks. Each of the three species included here started with similar resilience values (see Table 1). This is useful, as it gives us a common starting point to observe the change in resilience following the introduction of new nodes.

We computed prospective resilience under a number of different scenarios in order to determine the conditions under which networks would have the highest prospective resilience (i.e., which attachment mechanism is the most effective for maximizing thee network’s prospective resilience). In each condition, we calculate the prospective resilience by adding 20 new nodes to each network. We varied the number of new links, *m*, that each new node added to the network (*m* = 4, 8 and 16). Each simulation was repeated 100 times and the means and standard deviations were recorded from these runs. The resilience was calculated with a rate of node removal, *b* = 50 (see Section 3.1).

The results comparing the prospective resilience across the three species and attachment mechanisms are shown in Figure 4. We found that the most effective mechanism for adding new nodes to the networks was the attachment rule based on the *gene expression* of nodes in the original network.

**Figure 4:**
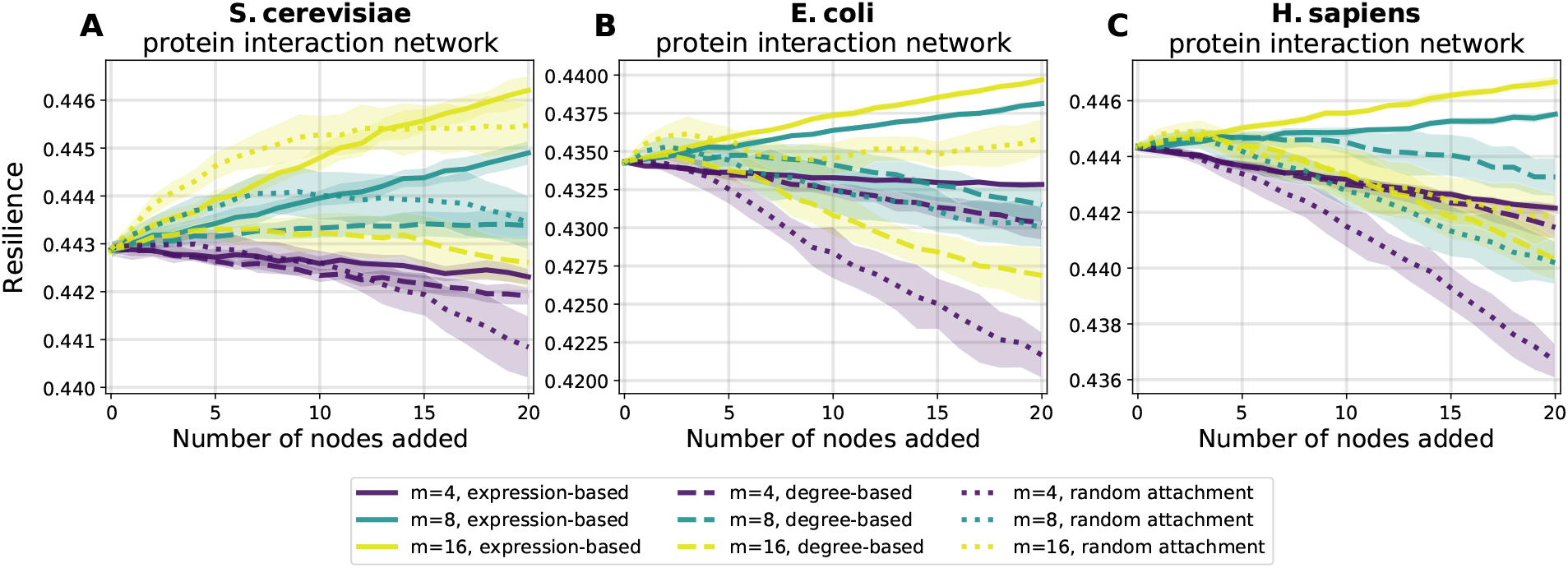
Prospective resilience of three model ribosomal networks. As more nodes are added (horizontal axes), the resilience of the resulting network changes (vertical axes). The colour of each curve corresponds to the number of new links that each new node enters the network with, and the line style (solid, dashed, or dotted) corresponds to the three different node-attachment mechanisms, as indicated in the legend.

Degree-based and random attachment were on average less effective at increasing the resilience of these networks. In general, a higher positive slope indicated that the attachment rule (along with the number of links that each new node enters the network with) generated higher prospective resilience. For information about the statistical differences between the slopes of each curve in Figure 4, see SI 5.4.

In order to put these results in a better context, we performed a survey of resilience in random networks as the inference of network resilience has been under-explored for random networks. In SI 6.1, we include several explanatory simulations that offer a more comprehensive intuition about how this measure behaves in networks. We highlight two main behaviors of this measure: its dependence on the network density and the *degree heterogeneity* of the network. We illustrated this further in the context of Erdős-Rényi networks (SI 6.1.1) and preferential attachment networks (SI 6.1.2).

Based on our analyses of random networks, adding more links (therefore making the resulting network more dense) increased the prospective resilience in each of the three networks. This is shown by the different colored lines in Figure 4. This holds regardless of the method of attachment. In other words, given that *links* in these networks correspond to interactions between proteins, our results suggest that a network’s resilience is more likely to increase if novel proteins are highly interactive and particularly if they are highly interactive with highly expressed proteins that are already present in the network.

#### 3.2.2 Resilience and modularity

We found that the gene expression-based attachment mechanism was most effective at maximizing the prospective resilience of the three networks included here. This finding does not immediately account for the extent to which this could have been due to higher-order, structural (i.e., not necessarily biological) properties of the network. To address this, we tested whether the observed results could be explained by other network properties—in this case, the modularity. In general, we refer to networks as being modular when they consist of densely-connected clusters of nodes that are connect more to each other than to the rest of the network. We chose to analyse modularity due to observations of strong modular structures in the ribosomal networks, especially in the case of *H. sapiens* (Figure 2). Additionally, we note that the three networks have very different initial levels of modularity (Table 1).

Here, we examine whether we observe similar results to those in Section 3.2.1 if we instead look at the change in the networks’ modularity following the introduction of new nodes. To do this, we computed the modularity of the network after each addition of new nodes. Full details of the analysis are found in Section 5.3.1. We found that the behavior of prospective modularity did not resemble the observed trends for prospective resilience (Figure 5). In fact, node addition affected the prospective modularity of each network differently, with no discernible pattern between the different networks. As such, modularity was ruled out as an explanatory measure for network resilience. In conclusion, the modular structure of the networks included here did not drive their prospective resilience.

**Figure 5:**
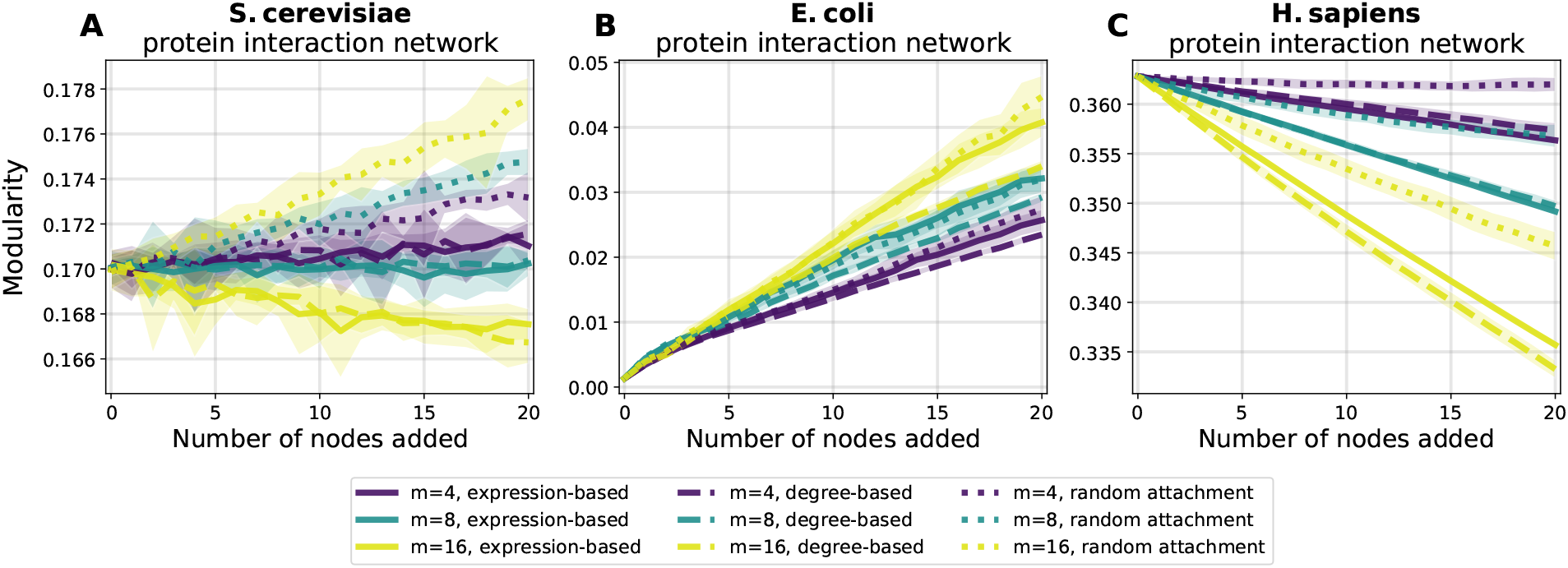
Prospective modularity of three model ribosomal networks. As a comparison measure, we also examined how the modularity of the network changes following the addition of new nodes. The colour scheme and line styles are the same as in Figure 4. Crucially, we do not find any evidence that the prospective resilience results observed in Figure 4 are being driven by the change in the networks’ community structures, as the three plots above show highly divergent patterns, suggesting that there is a more distinct mechanism underlying the prospective resilience.

#### 3.2.3 Noise and protein networks

We previously observed that gene expression was moderately correlated with node degree while gene expression-based attachment performed better than degree-based attachment. Here, we examine how decoupling of gene expression from the network topology affects the prospective resilience of the network. In other words, we probe to what extent the performance of gene expression-based attachment is influenced by the distribution (i.e. Figure 2D, 2E, and 2F) of gene expression values and its potential to create novel network structure, rather than any relationship between the gene expression values and the PPI network’s existing topology. To do this, we randomly shuffled the gene expression values across the network and re-ran the prospective resilience simulations. We did this for different amounts of shuffling. For example, at 20% shuffling, the gene expression values for a randomly chosen 20% of the proteins (network nodes) were subject to a random permutation, while the remaining 80% of proteins retained their original gene expression. At 100% noise the gene expression values were randomly assigned to nodes across the network.

We observe, in each of the three networks, that elevated shuffling of gene expression *increased* prospective resilience (Figure 6). In other words, biological noise simulated as random distribution of expression, increases prospective resilience. It makes sense that some noise would increase the prospective resilience; resilience increases as networks becomes more dense, and shuffling the gene expression values may increase the chance that a given node receives a link from an incoming node. However, increasing noise always increased the prospective resilience. This can be explained by how our computational simulation does not regard limitations or consequences regarding biological functionality, merely protein interaction and PPI network structure.

**Figure 6:**
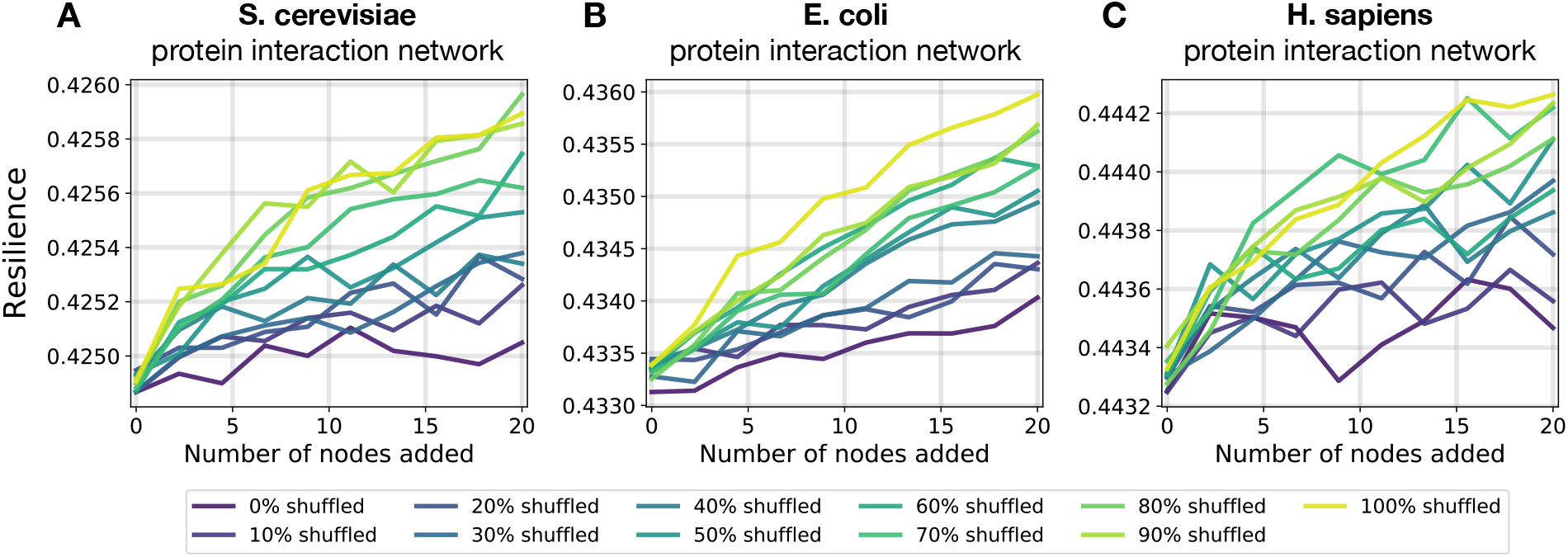
Prospective resilience and randomized gene expression. Here, we ask whether the specific gene expression of the proteins in these three networks is driving the high prospective resilience of the expression-based attachment rule or whether merely attaching based on a shuffled gene expression distribution could bring about these results. In each of the three panels above, we see that the prospective resilience of the networks *increases* simply by increasing the fraction of nodes with shuffled gene expressions. Note: for the three panels above, each new node joins with *m* = 5 for *S. cerevisiae* and *E. coli*, and *m* = 6 for *H. sapiens*. These values were selected so that the slope of the prospective resilience would be closest to 0.0 when the gene expression was not shuffled (0% shuffled). See Table 2 for how the correlation between a node’s degree and its gene expression changes as noise increases.

Therefore, we conclude that the effect of the uneven distribution of gene expression (and its limited association with degree) on the preferential attachment mechanism promotes new hubs (higher degree nodes) of connectivity in the network, which increases the network’s prospective resilience. The greater the novelty in the network structure created by this mechanism (i.e. the less correlation between degree and gene expression) the greater the network’s prospective resilience.

## 4 Discussion

This study used new network scientific methods to undertake a systems approach to understanding how novelty is incorporated into protein-protein interaction networks. We accomplished this by adapting a measure of *network resilience* to characterize the *prospective resilience* of three ribosomal protein networks. We found that the prospective resilience of the *S. cerevisiae*, *E. coli*, and *H. sapiens* ribosomal networks was greatest when node addition was based on the gene expression of the proteins in the original networks. This suggests that the distributed levels of gene expression among proteins facilitates or enables the system of interacting proteins to receive and incorporate new proteins. It also suggests an important correspondence between the structure and biological properties of protein networks.

We also undertook a comprehensive survey of how network resilience behaves in random and preferential attachment networks, and highlighted its dependence on the density and degree heterogeneity of the network (SI 6.1). These simulations contextualize the analyses that we performed for ribosomal networks and provide a platform for further use of the metric in a more theoretical sense.

We compared the prospective resilience to a meso-scale network structural measure (which we refer to as the *prospective modularity*) to determine if the observed increases in resilience was due to the more widely studied property of community structure [37]. No clear trend between prospective resilience and prospective modularity was found between the networks (Figure 5). This supports the hypothesis that there remains a crucial role of gene expression specifically in the resilience of a PPI network.

In a biological setting, network resilience infers biological redundancy. We assume that novel proteins can be integrated into existing PPI networks if they do not cause dis-connectivity of the network, and instead add to the network redundancy. In other words, we assume that novel proteins are likely to be integrated into an existing PPI network if they elevate the network redundancy. We find that likelihood of a novel protein being integrated is dependent on the existing topology of PPI and internal connectivity, but also gene expression. The results of our node attachment analysis imply that novel proteins are able to be integrated if they i) are interactive with many existing proteins, or ii) primarily interact with proteins that are already highly interactive and abundant (inferred by gene expression) [38].

Our findings suggest that novel proteins might enter PPI networks and interacting broadly as generalists. This is in line with previous research that suggests how many proteins, i.e. enzymes, begin as generalists with many interacting partners, and later evolve more specialized interactions [38, 39]. We speculate that novel proteins may be conserved prior to gaining a so called ‘important’ function, simply by being tolerated and adding to the network resilience, as suggested in research on *de novo* genes [6, 7]. *De novo* genes are found to have both high, stable and stochastic gene expression [40]. Future research should address to what extent gene expression enables *de novo* genes to integrate in PPI networks, by comparing the topologies of highly conserved PPI networks with PPI networks that have undergone evolutionary recent topological alteration, e.g. where *de novo* genes are integrated. However, this should not only be limited to analyses of *de novo* genes, but any protein acquiring a novel function (e.g. new interaction partner or catalytic function).

The results from the noise analysis, Section 3.2.3, suggest that shuffling gene expression tends to further increase resilience. The tails of the gene expression distributions may indicate that i) the most important factor for increasing resilience is the creation of new hubs of connectivity (new nodes strongly connecting to a few existing nodes), and ii) these new hubs are more effective in increasing resilience if created randomly in the network and not correlated with the already established topology. Interestingly, a heavy-tailed (log-normal) factor of attachment has been recently demonstrated as an accurate explanation of the degree distributions across various complex networks [41], lending credence to the idea of gene expression as (at least part of) such an explanatory mechanism in PPI networks. If gene expression influences the evolution of the PPI network, then it necessarily needs to have an amount of correlation with the existing degree distribution of the network. Thus, even though we observe that the completely randomised gene expression across the network yields a more resilient network, given enough time, the network connectivity would evolve to correlate with the new gene expression values of the corresponding proteins. Then, more noise would be required to increase the network resilience. In an biological setting, we assume resilience to be important for functional performance, but not more important than the biological function of the network. In an evolutionary trajectory of a PPI network, we would thus expect to see a trade-off between the topological influence of gene expression (i.e. correlation between gene expression and protein node degree) and the emergence of novelty through biological noise (i.e. weakened correlation between node degree and gene expression). Arguably, this is reflected in the weak to moderately strong correlations found in Figure 2G, 2H, and 2I. This conforms to classic theoretical notions of the usefulness of noise in biological systems [42, 43]. Further research is needed to determine the extent to which this holds for biological networks in general.

Subsequent and systematic analyses of the prospective resilience of other species’ ribosomal networks (not to mention gene pathway networks, metabolic networks, etc.) will allow researchers to form more precise hypotheses about other possible mechanisms—especially ones relating gene expression, pair-wise protein interactions and overall PPI network topology—which might be driving the results we observe and delineate here. In addition, it would be useful to explore how prospective resilience changes under other biologically-informed methods for introducing proteins into PPI networks. For example, the measured interaction strength between proteins is used to define presence or absence of interaction here but it may also be used to create a weighted network. Novel proteins, e.g. duplicates of existing proteins, may have their attachment probabilities formed based on the interaction strength that the original protein has with other proteins. However, we view this work as a first step towards understanding the stability of a network’s resilience to novel information. Moreover, prospective resilience is a measure that can describe networks in general; it is particularly meaningful in the study of biological systems, but since complex systems are often described as recapitulating common properties across different domains, this network measure can be used in any system that undergoes and incorporates novel information.

## 5 Methods

### 5.1 Data sources

We make use of publicly available data of protein interaction networks from Zitnik et al. (2019). Full interactomes were obtained from their website (SNAP) for 3 model organisms: *Saccharomyces cerevisiae*, *Homo sapiens*, and *Escherichia coli* str. K12 [28]. We additionally gathered gene expression data for each of the species studied. Expression data for *S. cerevisiae* came from the wildtype data accessible on the NCBI GEO database (accession: GSE67387) [32, 33]. The GTEx Consortium [35] collected *H. sapiens* gene expression data for various tissues, which was accessed via the EMBL-EBI Expression Atlas [34]. We utilized expression reported in the spleen as it was the tissue where most of the genes in the ribosomal network were expressed. Wildtype gene expression data for *Escherichia coli* str. K12 substr. MG1655 (NCBI:txid511145) was obtained from the NCBI GEO database (accession: GSE48829) [32, 36]. Meysman et al. (2013) originally reported expression as count data; we converted from counts to transcripts per million (TPM) with custom R scripts and gene lengths for *Escherichia coli* str. K12 retrieved from UniProt [44] in June 2019. To convert to TPM, we first divided the read counts by the length of each gene (in kilobases) to get reads per kilobase (RPK). The sum of all RPK values was divided by one million to produce a scaling factor, which was then multiplied by each protein’s RPK to produce their expression in TPM.

### 5.2 Network resilience

A network, *G*, consists of *N* nodes, *V* = {*υ*_1_, *υ*_2_, …, *υ_N_*}, connected by M links, *E* = {(*υ_i_*, *υ_j_*) : *υ_i_*, *υ_j_* ∈ *V*}. The resilience of a network is based on an information theoretic analysis of the distribution of the sizes of connected components in *G* [20]. A connected component may be defined as follows. If there exists a path of links between two nodes, *υ_i_* and *υ_j_*, in *G*, then they are in the same connected component, *c_x_*, of *G*. Otherwise *υ_i_* and *υ_j_* are in separate components, *c_x_* and *c_y_*, say, of *G*. If *υ_i_* has no links, and thus no paths from itself to any other node in *G*, then *υ_i_* is an isolated component of *G*. From this, we see that *G* is composed of *X* disjoint connected components, 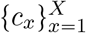, of varying sizes such that 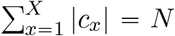. We can then confer a notion of probability to each component proportional to its size, *p_x_* = |*c_x_*|/*N*, such that if we chose a node at random from *G* it would have probability *p_x_* of coming from component *c_x_*. Resilience is then measured through a modified Shannon diversity of the connected component size distribution in the presence of node removal [20], as follows:

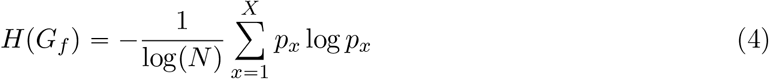

This value is minimal, *H*(*G_f_*) = 0, when the network consists of a single connected component where paths exist between all node pairs, since log1 = 0, and maximal, *H*(*G_f_*) = 1, when the network consists only of isolated components—*H*(*G*) = −log(*N*^−1^)/log *N* = 1. Through simulating the removal of a fraction of randomly-selected nodes, *f*, in a given network by removing all links to those nodes and leaving them as isolated components, we are left with a new network, *G_f_*. Then the entropy of the connected component distribution will increase with increasing *f*. With an increasing fraction of randomly-removed nodes, *f*, the entropy of the number of connected components will increase until *f* = 1.0, at which point there are *N* disconnected nodes (isolated components), reducing the network to the maximal case of *H*, as previously noted. We show an example of this process, as *f* increases, for an arbitrary simulated network (Figure 1). The resilience, *R*(*G*) of a network, *G*, is then defined as follows:

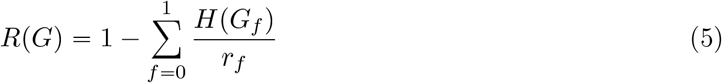

where *r_f_* is the rate of node removal such that 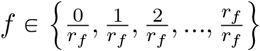. In this work, we default to a value of *r_f_* = 100, which means that the calculation of a network’s resilience involves iteratively removing 0%, 1%, 2%, …, 100% of the nodes in the network. For each value of *f*, we simulate the node removal process 20 times.

### 5.3 Structural modularity measure

#### 5.3.1 Network modularity

Networks are often analyzed by their community structure—that is, to what extent do nodes in a network connect to other similar nodes, whether in their structural properties or specific attributes [37, 45–48]. There are a number of different ways to detect community structure in networks, from algorithmic optimization to statistical/inferential to dynamical approaches [45, 49, 50] (e.g., the color of the nodes in the networks in Figure 2 was determined by one such approach [48]). Regardless of the community detection approach, each method outputs a partition that maps each node to a given community. The *modularity* of a given partition is a number that scores the extent to which it captures nodes’ tendencies to connect to other nodes in their same community at the expense of nodes in other communities [37]. While imperfect, this measure endows us with a powerful intuition for assessing higher-order network properties; namely, a network with high modularity partitions is likely to have obvious clusters of nodes, structurally separated from other parts of the network.

#### 5.3.2 Prospective modularity

Here, we use the notion of modularity in an attempt to give possible explanations for the network mechanisms behind the observed trends in the prospective resilience of the ribosomal networks studied in this work. In particular, we define *prospective modularity* in the same vein as our prospective resilience measure to compare how node addition impacts resilience and modularity. The prospective modularity (*PM*) of a network is defined as the change in modularity following the addition of new nodes to a network (note the precise similarities between this measure and the prospective resilience). The addition of a new node, *υ*_*t*+1_ with *m* disconnected links, to a network, *G_t_*, at time, *t* + 1 will likely change the modularity of the network. More specifically, by re-running a community detection algorithm on the resulting network, *G*_*t*+_ι, and calculating the modularity of the resulting partition, we can observe the stability of this partition over time and ask whether the modularity will increase or decrease. Further, by varying the node-addition mechanism (adding nodes randomly, preferentially based on degree, or preferentially based on gene expression), we can observe the different effects that network structure and gene expression has on the prospective modularity of a given network.

### 5.4 Statistics of prospective resilience and modularity

In order to determine the extent to which the curves in Figure 4 differ from one another, we perform a series of statistical tests. The curves represent the average of 10 independent simulations for each condition. We utilize all existing simulation data here. For each value of *m* in each species, we perform an ANCOVA for each pair of attachment methods. We do a Bonferri-correction to correct for multiple testing and obtain a significance cutoff at *p* = 0.0166. Additionally, we calculate Cohen’s *d* from the *F*-statistic presented by the ANCOVA. The p-values and effect size (Cohen’s d) for each comparison are presented in Table 3. Almost all of these slope comparisons are statistically significant. We do the same pairwise ANCOVA and effect size comparisons for the curves in Figure 5 and report the outputs in Table 4. For *S. cerevisiae*, *E. coli* and *H. sapiens*, the majority of slopes are significantly different and show significant differences for larger values of *m*.

**Table 3:** ANCOVA results for pairwise prospective resilience slope comparisons in Figure 4. Almost all comparisons are significant under the Bonferroni-corrected significance threshold (*p* < 0.0166). Cohen’s *d* was calculated from the ANCOVA’s *F*-statistic for each comparison.

| Species | m | Pairwise comparison | p-value | Cohen's d |
| --- | --- | --- | --- | --- |
| *S. cerevisiae* | 4 | expression-based, degree-based | $4.86e^{-16}$ | 1.31 |
| *S. cerevisiae* | 4 | expression-based, random attachment | $4.02e^{-11}$ | 0.89 |
| *S. cerevisiae* | 4 | degree-based, random attachment | $1.79e^{-07}$ | 0.62 |
| *S. cerevisiae* | 8 | expression-based, degree-based | $6.66e^{-26}$ | 2.55 |
| *S. cerevisiae* | 8 | expression-based, random attachment | $1.03e^{-08}$ | 0.71 |
| *S. cerevisiae* | 8 | degree-based, random attachment | $7.21e^{-01}$ | 0.04 |
| *S. cerevisiae* | 16 | expression-based, degree-based | $2.03e^{-25}$ | 2.47 |
| *S. cerevisiae* | 16 | expression-based, random attachment | $2.20e^{-04}$ | 0.40 |
| *S. cerevisiae* | 16 | degree-based, random attachment | $6.84e^{-10}$ | 0.80 |
| *E. coli* | 4 | expression-based, degree-based | $5.68e^{-27}$ | 2.73 |
| *E. coli* | 4 | expression-based, random attachment | $4.52e^{-33}$ | 4.00 |
| *E. coli* | 4 | degree-based, random attachment | $2.46e^{-29}$ | 3.17 |
| *E. coli* | 8 | expression-based, degree-based | $2.93e^{-23}$ | 2.15 |
| *E. coli* | 8 | expression-based, random attachment | $6.94e^{-30}$ | 3.28 |
| *E. coli* | 8 | degree-based, random attachment | $2.83e^{-06}$ | 0.54 |
| *E. coli* | 16 | expression-based, degree-based | $2.34e^{-33}$ | 4.07 |
| *E. coli* | 16 | expression-based, random attachment | $1.81e^{-15}$ | 1.26 |
| *E. coli* | 16 | degree-based, random attachment | $1.53e^{-19}$ | 1.68 |
| *H. sapiens* | 4 | expression-based, degree-based | $4.75e^{-12}$ | 0.96 |
| *H. sapiens* | 4 | expression-based, random attachment | $1.37e^{-22}$ | 2.06 |
| *H. sapiens* | 4 | degree-based, random attachment | $6.30e^{-21}$ | 1.85 |
| *H. sapiens* | 8 | expression-based, degree-based | $1.90e^{-15}$ | 1.26 |
| *H. sapiens* | 8 | expression-based, random attachment | $1.39e^{-28}$ | 3.02 |
| *H. sapiens* | 8 | degree-based, random attachment | $1.72e^{-16}$ | 1.36 |
| *H. sapiens* | 16 | expression-based, degree-based | $1.12e^{-25}$ | 2.52 |
| *H. sapiens* | 16 | expression-based, random attachment | $1.56e^{-31}$ | 3.64 |
| *H. sapiens* | 16 | degree-based, random attachment | $2.74e^{-04}$ | 0.39 |

**Table 4:** ANCOVA results for pairwise prospective modularity slope comparisons in Figure 5. Many comparisons are significant based on the Bonferroni-corrected significance threshold (*p* < 0.0166). Cohen’s *d* was calculated from the ANCOVA’s *F*-statistic for each comparison.

| Species | m | Pairwise comparison | p-value | Cohen's d |
| --- | --- | --- | --- | --- |
| *S. cerevisiae* | 4 | expression-based, degree-based | $6.93e^{-01}$ | 0.04 |
| *S. cerevisiae* | 4 | expression-based, random attachment | $6.65e^{-14}$ | 1.12 |
| *S. cerevisiae* | 4 | degree-based, random attachment | $4.07e^{-12}$ | 0.97 |
| *S. cerevisiae* | 8 | expression-based, degree-based | $4.49e^{-01}$ | 0.07 |
| *S. cerevisiae* | 8 | expression-based, random attachment | $4.71e^{-25}$ | 2.42 |
| *S. cerevisiae* | 8 | degree-based, random attachment | $4.07e^{-25}$ | 2.43 |
| *S. cerevisiae* | 16 | expression-based, degree-based | $2.34e^{-01}$ | 0.12 |
| *S. cerevisiae* | 16 | expression-based, random attachment | $6.31e^{-36}$ | 4.77 |
| *S. cerevisiae* | 16 | degree-based, random attachment | $2.46e^{-33}$ | 4.07 |
| *E. coli* | 4 | expression-based, degree-based | $7.63e^{-06}$ | 0.51 |
| *E. coli* | 4 | expression-based, random attachment | $3.24e^{-04}$ | 0.39 |
| *E. coli* | 4 | degree-based, random attachment | $4.13e^{-11}$ | 0.89 |
| *E. coli* | 8 | expression-based, degree-based | $4.13e^{-06}$ | 0.52 |
| *E. coli* | 8 | expression-based, random attachment | $1.92e^{-01}$ | 0.13 |
| *E. coli* | 8 | degree-based, random attachment | $1.32e^{-05}$ | 0.49 |
| *E. coli* | 16 | expression-based, degree-based | $8.36e^{-13}$ | 1.03 |
| *E. coli* | 16 | expression-based, random attachment | $1.06e^{-04}$ | 0.42 |
| *E. coli* | 16 | degree-based, random attachment | $1.74e^{-16}$ | 1.36 |
| *H. sapiens* | 4 | expression-based, degree-based | $3.00e^{-24}$ | 2.30 |
| *H. sapiens* | 4 | expression-based, random attachment | $6.39e^{-36}$ | 4.77 |
| *H. sapiens* | 4 | degree-based, random attachment | $1.05e^{-32}$ | 3.91 |
| *H. sapiens* | 8 | expression-based, degree-based | $4.65e^{-08}$ | 0.66 |
| *H. sapiens* | 8 | expression-based, random attachment | $5.60e^{-30}$ | 3.30 |
| *H. sapiens* | 8 | degree-based, random attachment | $1.49e^{-27}$ | 2.83 |
| *H. sapiens* | 16 | expression-based, degree-based | $1.80e^{-10}$ | 0.84 |
| *H. sapiens* | 16 | expression-based, random attachment | $4.43e^{-35}$ | 4.53 |
| *H. sapiens* | 16 | degree-based, random attachment | $4.90e^{-33}$ | 3.99 |

## Acknowledgements

This work was conceived at and supported by the Santa Fe Institute (SFI) Complex Systems Summer School (CSSS) in June, 2019 and a working group grant in December, 2019. The authors thank Douglas Reckamp for his contribution to early discussions about networks and novelty. B.K. is supported by the National Defense Science & Engineering Graduate Fellowship (NDSEG). A.S. acknowledges support from NSF award DGE-1632976. M.M.J. acknowledges support from NIH training grant 1T32LM012414-01A1. K.M.S. acknowledges support from Health Data Research UK, an initiative funded by UK Research and Innovation Councils, National Institute for Health Research (England) and the UK devolved administrations, and leading medical research charities. A.I.T is supported by the SFI. A.S.K acknowledges support from the Independent Research Fund Denmark (DFF-4181-00490), and support from Studienstiftung des deutschen Volkes (DE) during the time when this project was initially conceived.

## Author contributions statement

A.S.K. conceived the project with B.K. All authors contributed to the analyses in this work. Data were retrieved by A.S.K. and M.M.J. All authors contributed to the writing of the article.

## Data availability and software

Data, software, and reproducibility materials—including Python code with examples—can be found at https://github.com/jkbren/presilience.

## 6 Supplementary Information

This supplementary information consists of theoretical and experimental work on the general properties of network resilience (Section 6.1), and supplementary Tables 3 and 4 on statistical results on differences of the network prospective resilience and modularity curves for different attachment mechanisms.

### 6.1 Resilience in random networks

Here, we describe resilience as calculated on two different random network models. One set of networks is generated by the Erdős-Rényi model and the other set is composed of preferential attachment models. Calculating the resilience of these networks offers insight into the behaviour of this resilience measure. In addition, we show here that the theoretical upper bound of the resilience measure is 0.5.

#### 6.1.1 Resilience in Erdős–Rényi networks

In Erdős–Rényi networks nodes are connected uniformly at random. That is, each new node has a probability *p* to connect to one of the *N* nodes present in the network. In this way, the parameter *p* dictates the density of the network since a higher *p* means that each new node is more likely to form more edges (more of the possible edges between nodes that can exist, exists). Plotting resilience against *p* will then show us something about the relationship between resilience and density. Indeed, we observe a positive relationship between density and resilience (Figure 7 right). For a network with density close to 0 (generated by a very low *p*) the resilience is also close to 0 and conversely a complete graph *p* = 1 yields a high resilience.

**Figure 7:**
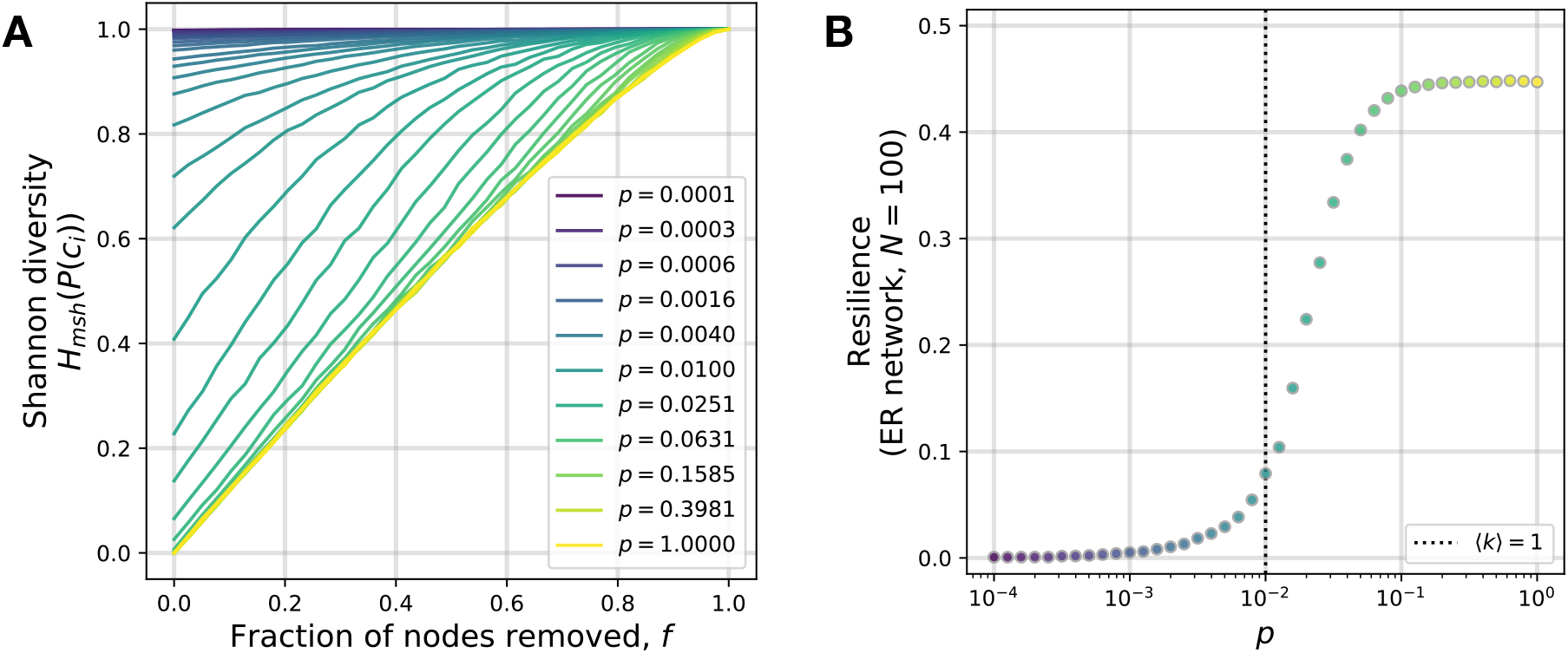
Erdős–Rényi and resilience. Shannon Diversity changes as nodes are removed from random attachment networks of 100 nodes for different values of *p* **(A)**. For *p* < 0.4, the networks tend to have disconnected components before any of the nodes are removed. This means that the Shannon diversity is greater than zero before any of the nodes are removed. In addition, the resilience values for the preferential attachment networks corresponding to **(A)** are shown in **(B)**.

#### 6.1.2 Resilience in preferential attachment networks

Preferential attachment networks are generated by the addition of new nodes, each with *m* dangling links (or disconnected links) [51, 52]. These *m* links connect to nodes, *υ_j_*, that are already present in the network based on a probability proportional to 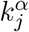. Where *k_j_* is the degree of node *υ_j_* and *α* a parameter which gives the amount of preferential attachment. *α* = 0 means that there is no preferential attachment and the probability of attaching a dangling link to a node is the same for all nodes. *α* > 0 means that dangling links are more likely to attach to nodes that have a high degree (nodes that already have a lot of links attached to them) and means that there is positive preferential attachment. Finally, *α* < 0 means that dangling links are more likely to attach to nodes that have a low degree.

We observed that the resilience of a preferential attachment network depended on both *α* (Figures 8 B, D & F) and *m* (which varies between rows of subfigures in Figure 8). Looking at the network where the number of dangling links, *m*, is 1, (Figure 8B), we see that resilience varies to a large extent as the tuning parameter *α* varies (roughly between 0.19 and 0.32). As *m* increases the relationship between *α* and resilience changes and now low values of *α* yield higher values of resilience. In addition the spread of resilience decreases drastically, already for *m* = 2 resilience varies roughly only between 0.38 and 0.39 for different values of *α*. For *m* = 24, resilience hardly varies at all and there is no clear relationship between *α* and resilience. A higher *m* means that the networks are more dense. Therefore it seems that for dense networks, the structure of the network (governed by *α*) plays a less important role.

**Figure 8:**
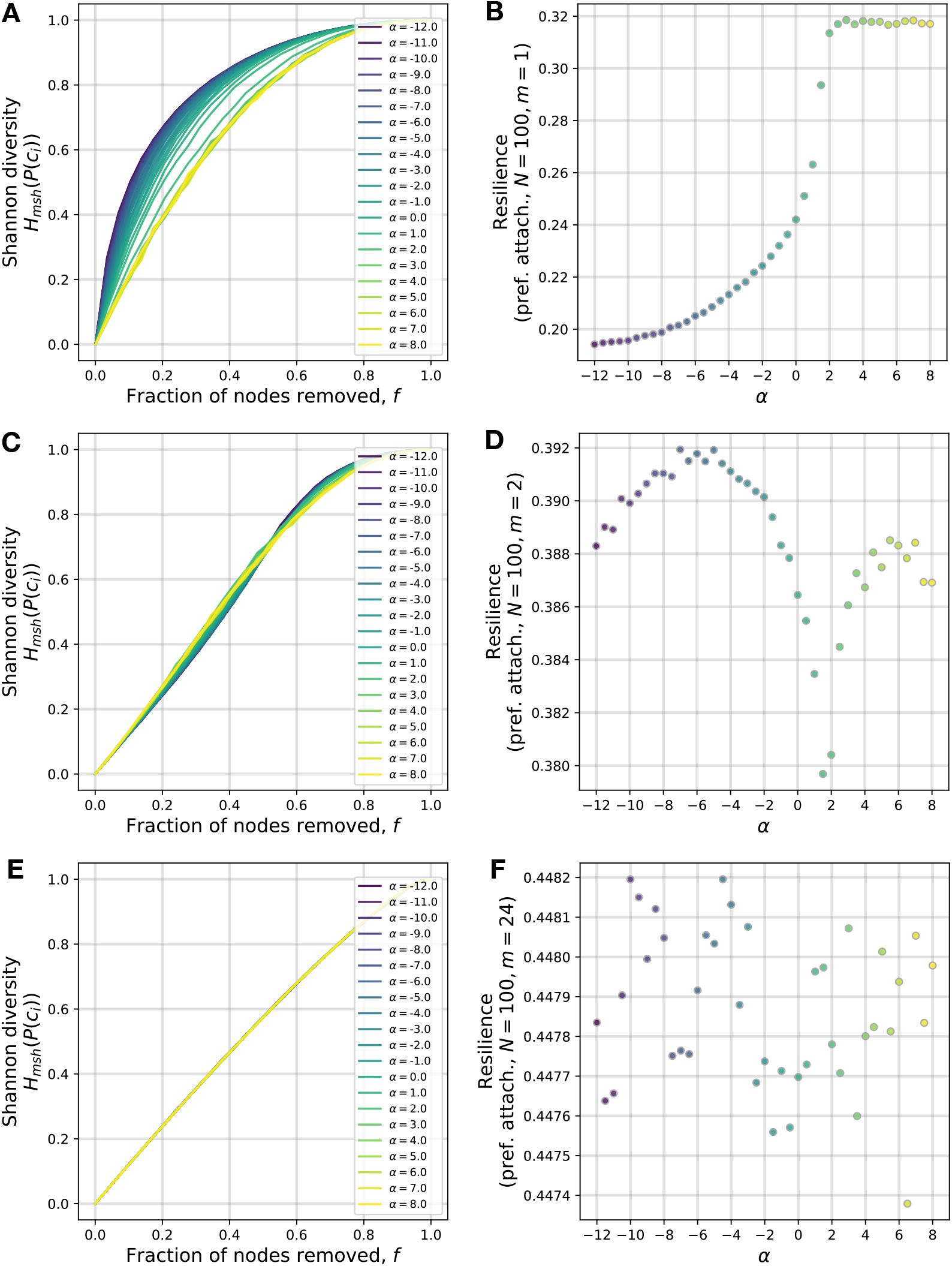
Preferential attachment and resilience. Shannon Diversity changes as nodes are removed from random attachment networks of 100 nodes for different values of *α* (Figures 8A, 8C, and 8E). The resilience values for the preferential attachment networks corresponding to the left plot are shown in Figures 8B, 8D, and 8F. The difference between the rows of plots are the number of dangling edges, *m*, a node has as it enters the network when the network is being generated.

#### 6.1.3 Upper bound for network resilience

It was originally claimed that resilience takes values in [0, 1] [20]. However, we show that the upper bound of resilience is not higher than 0.5 as this is achieved asymptotically by the continuous counterpart of resilience applied to the complete graph of size *N* as *N* → ∞.

The maximum resilience for a network of size *N* is achieved if, when all the links adjacent to any *k* nodes are removed, we end up with those *k* nodes as isolated nodes and the rest of the network remains connected as one component of size *N* – *k*, for all *k*. A network which satisfies this property is the complete graph, *K_N_*. Take away the links of any *k* nodes in the complete graph and we have *k* isolated nodes and a complete subgraph of size *N* – *k*. In this instance, the modified Shannon diversity is:

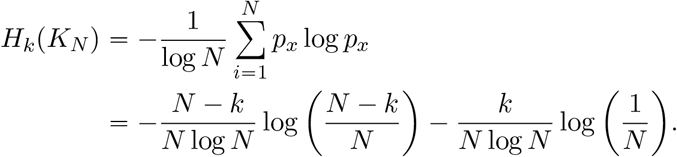

Then, taking the above as a continuous function, integrating between 0 and *N* with respect to *k* and dividing by *N* (i.e. taking the average of the function) we get:

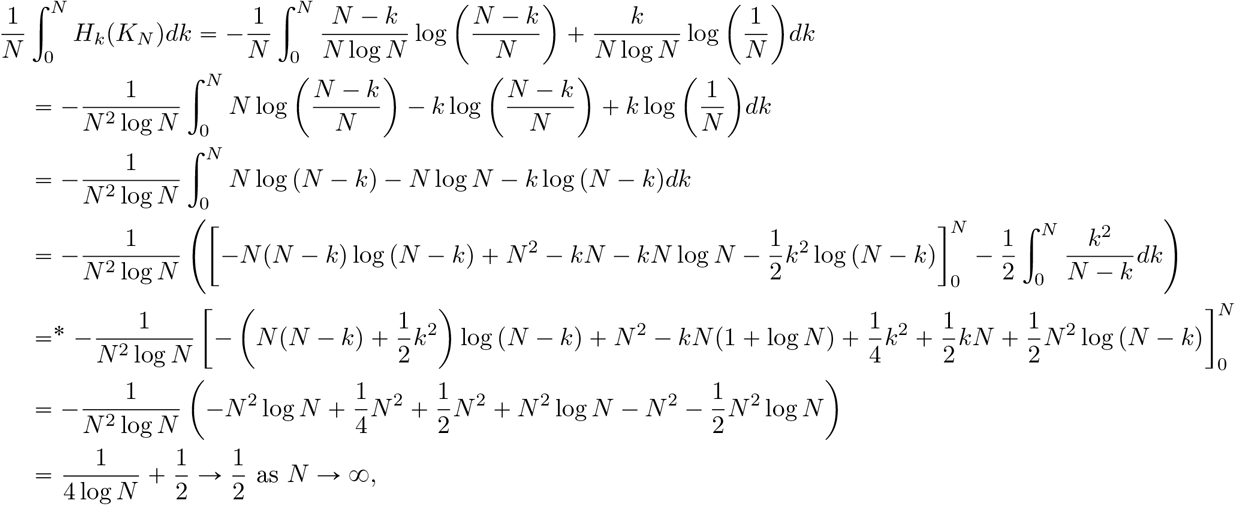

where * is achieved by polynomial division and then integration of the remaining integrand. Also, this is a decreasing function with respect to *N*, so 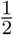 is the minimum of this integral with respect to *N*, so an upper bound of the continuous resilience function, *R_c_*(*G*) is given by 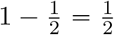. Since the discrete function, *R*(*G*) takes values equally spaced on *R_c_*(*G*) and *R_c_*(*G*) is a concave function, 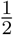 is also an upper bound for *R*(*G*).

